# Detection and Molecular Characterization of Rotavirus and Picobirnavirus in Wild Avians from Amazon Forest

**DOI:** 10.1101/2020.09.15.297689

**Authors:** José Wandilson Barboza Duarte Júnior, Elaine Hellen Nunes Chagas, Ana Carolina Silva Serra, Lizandra Caroline dos Santos Souto, Edvaldo Tavares da Penha Júnior, Renato da Silva Bandeira, Ricardo José de Paula Souza e Guimarães, Hanna Gabriela da Silva Oliveira, Thaymes Kiara Santos Sousa, Cinthia Távora de Albuquerque Lopes, Sheyla Farhayldes Souza Domingues, Yashpal Singh Malik, Felipe Masiero Salvarani, Joana D’Arc Pereira Mascarenhas

**Affiliations:** Evandro Chagas Institute, Ministry of Health, Ananindeua, Pará, Brazil; University of the State of Pará, Institute of Veterinary Medicine, Castanhal, Pará, Brazil; Indian Veterinary Research Institute, India

**Author notes:** Corresponding author: José Wandilson Barboza Duarte Júnior, Virology Section, Evandro Chagas Institute, Health Surveillance Secretariat, Ministry of Health, Rodovia BR 316-KM 07, S / N, Levilândia, 67.030-000, Ananindeua, Pará, Brazil.

**Keywords:** Genotyping, Genogroup I, Picobirnavirus, Rotavirus, Sequence analysis, Wild birds

## Abstract

The present study reports the detection and molecular characterization of rotavirus A (RVA), rotavirus D (RVD), rotavirus F (RVF), rotavirus G (RVG) and picobirnavirus (PBV) in fecal specimens of wild and exotic birds (n = 23) from different cities of Pará state, which were hospitalized at Veterinary Hospital of the Federal University of Pará, Brazil, between January 2018 to June 2019. The animals exhibited different clinical signs, such as diarrhea, malnutrition, dehydration and fractures. The results showed 39.1% (9/23) of positivity for RVA by RT-qPCR. Among these, one sample (1/9) for the NSP3 gene of T2 genotype was characterized. About 88.9% (8/9) for the VP7 gene belonging to G1, equine-like G3 and G6 genotypes, and 55.5% (5/9) for the VP4 gene of P[2] genotype were obtained. In the current study, approximately 4.5% of the samples (1/23) revealed coinfection for the RVA, RVD and RVF groups. Furthermore, picobirnavirus (PBV) was detected in 1 of the 23 samples tested and was classified in the Genogroup I. The findings represent the first report of the circulation of RVA, RVD, RVF, RVG and PBV genotypes in wild birds in Brazil and suggest the possible interspecies transmission of RVs and PBVs.

## INTRODUCTION

The Brazilian Amazon biome is the largest ecosystem of wildlife biodiversity, and Brazil is the third country that protects the largest diversity of birds in the world, with registration of 1.919 species [1] and global distribution of 10.429 bird species [2]. Pará is the second state with the largest territorial extension covering the Amazonian biome, but is also a region that suffers anthropic pressure with the advance of deforestation, fires, hunting and illegal sale of both wild and exotic species. This situation favors the proximity of wildlife with other animals and humans, which can lead to maximization and dispersion of zoonotic pathogens [3,4].

In wild ecosystems, several enteric agents may be present, including rotaviruses. Belonging to the family *Reoviridae*, genus *Rotavirus* is of paramount importance. Rotaviruses are icosahedral, with 11 segments of double-stranded RNA, no lipoprotein envelope and are classified in nine groups (A-I), based on the antigenicity of the VP6 protein. Accordingly, the groups A, D, F and G have been reported in avian species, with clinical manifestation or no manifestation, however, the circulation in wild birds is scarce in the Amazon region [5,6,7].

Picobirnavirus (PBV) belongs to the order *Diplornavirales*, family *Picobirnaviridae* and genus *Picobirnavirus* [8,9]. Picobirnaviruses have icosahedral symmetry, no lipoprotein envelope and genetic material consisting of 2 segments of double-stranded RNA [8]. The segment 1 encodes for two proteins, one whose function is still unknown and another one for the capsid protein. The segment 2 codifies for RdRp and allows the classification of PBV in Genogroup I (G-I) and Genogroup II (G-II), which have been reported in several species of animals, mainly in birds [8,10]. With regard to wild birds, Masachessi et al. [11] have previously detected the circulation of PBV in rhea, pheasant, pelican, Chinese goose and darwin-nandu (*Rhea pennata* or *Pterocnemia pennata*) in Argentina, but the occurrence of this virus in other species of wild and exotic birds is still unknown.

In this context, due to the limited knowledge on the epidemiology of these viruses and their impact in Brazilian wildlife, the present study herein reports the circulation of RVA, RVD, RVF and PBV in wild and exotic birds in Brazilian Amazon.

## METHODS

### Ethical aspects

The Biodiversity Authorization and Information System (SISBIO) of the Chico Mendes Institute for Biodiversity Conservation (ICMBio), Ministry of the Environment under No. 67300, and the Ethics Committee on the Use of Animals of the Evandro Chagas Institute (CEUA/IEC) under No. 03/2019 approved this research.

### Study area

From January 2018 to June 2019, 23 fecal samples were collected from different species of wild birds, including two samples from *Ramphastidae*, three from *Turdidae*, four from *Strigidae*, four from *Accipitridae*, one from *Falconidae*, one from *Fringillidae*, two from *Thraupidae*, one from *Cacatuidae* and five samples from *Psittacidae* that were hospitalized in the Veterinarian Hospital of the Federal University of Pará (HOVET-UFPA), in Castanhal, recognized as a reference in clinical care for wild animals in Pará state.

The diarrheic and non-diarrheic animals came from eight different cities of Pará: Belém, Benevides, Capanema, Capitão Poço, Castanhal, Inhangapi, Paragominas and Santa Izabel.

### Collection of clinical specimens

All fecal specimens were collected fresh, immediately after excretion, using sterile plastic bags, avoiding contact with contaminating materials. Then, all samples were labeled and stored in sterile, sealed and refrigerated universal collector tube at -20 °C.

### Laboratory methodology

Fecal suspensions were prepared at 10% (w/v) in phosphate buffered saline (PBS, 1X pH 7.4) and nucleic acid was extracted following the Boom et al. [12] method. Polyacrylamide gel electrophoresis (PAGE) and silver staining were applied in all specimens to detect rotavirus and picobirnavirus, according to Pereira et al. [13].

The detection of RVA was performed by RT-qPCR using the NSP3 gene as a target according described by Zeng et al. [14]. The samples with Cycle Threshold (CT) ≤ 40 were considered positive.

For RT-PCR/Nested reaction, cDNA was obtained by adding 4μL of dsRNA, 1μL of each primer pair 20mM, 2 μL of DNTP, 2.5μL of 5X buffer, 0.75μL of MgCl, 0.5μL of Reverse Transcriptase enzyme and 14.75μL of RNAse/DNAse free H_2_O for a final volume of 25μL. The cDNA was amplified to a final volume of 50μL, added to the 3μL of dNTP mix, 2.5μL of 5X buffer, 0.75μL of MgCl, 0.25μL of Taq Polymerase (Invitrogen^®^) and 18.50μL of RNAse/DNAse free H_2_O.

For the molecular characterization of RV, several primer sets were used (**Chart 1**) including: a) Gen-NSP3-F/R primers for the RVA NSP3 Gene [15]. b) N-VP7F1/R1 and N-VP7F2/R2 for the RVA VP7 gene [16]. c) N-VP4F1/R1 and N-VP4F2/R2 for the RVA VP4 gene [16]. d) SEG 10-C-S (+)/(-) for the RVA NSP4 gene [17]. e) RD6F/R, RF6F/R and RG6F/R for the RVD [18], RVF [18] and RVG [19], respectively, amplifying the VP6 genes.

**Chart 1.** Primers sets used to RV genotyping.

| NAME | GROUP | GENE | AMPLICON | METHOD |
| --- | --- | --- | --- | --- |
| Gen-NSP3-F/R | RVA | NSP3 | 1078 pb | RT-PCR<br>[15] |
| N-VP7F1/R1 | RVA | VP7 | 333 pb | RT-PCR<br>[16] |
| N-VP7F2/R2 | RVA | VP7 | 193 pb | NESTED<br>PCR [16] |
| N-VP4F1/R1 | RVA | VP4 | 257 pb | RT-PCR<br>[16] |
| N-VP4F2/R2 | RVA | VP4 | 214 pb | NESTED-<br>PCR [16] |
| SEG 10-C-S<br>(+)/(-) | RVA | NSP4 | 642 pb | RT-PCR<br>[17] |
| RD6F/R | RVD | VP6 | 742 pb | RT-PCR<br>[18] |
| RF6F/R | RVF | VP6 | 846 pb | RT-PCR<br>[19] |
| RG6F/R | RVG | VP6 | 876 pb | RT-PCR<br>[19] |

Chart1

For the detection of picobirnavirus, the primer pair PBV1.2 FP/RP was used according to the protocol described by Malik et al. [20]. In order to characterize the Genogroup I, nested PCR was performed using a primer pair designed by Dr. Yashpal S. Malik of Indian Veterinary Research Institute, India, Malik-2-FP forward (5’-TGG GWT GGC GWG GAC ARG ARGG-3’) and reverse Malik-2-RP (5’-YSC AYT ACA TCC TCC AC-3’) that amplifies a partial 580 bp fragment of RdRp.

The Sanger’s di-deoxy method of nucleotide sequencing [21] was performed. The sequences were aligned and edited using Geneious software v.10.0.6 [22] and then compared with other sequences deposited in GenBank through the Basic Local Alignment Search Tool (BLAST). Phylogenetic trees were constructed by MEGA V.6 program [23], based on Kimura parameters [24], using the non-parametric reliability test with bootstrap of 1000 replicas.

## RESULTS

The analysis by PAGE showed negativity for 23 samples, and there was no electrophoretic migration characteristic of PBV and RV. However, RT-qPCR was positive in six of eight cities for RVA, being detected in 39.1% (9/23) of avian fecal specimens from Benevides, Castanhal, Capanema, Inhapagi, Paragominas and Santa Izabel (**Chart 2**). No case was registered in Belém and Capitão-Poço. In turn, Castanhal concentrated 44.5% (4/9) of the cases and exhibited positivity for both RV and PBV (**Fig 1**).

**Chart 2.** Positive RV and PBV samples from animals hospitalized at the Veterinary Hospital of the Federal University of Pará, Brazil. **Legend:** Neg = Negative.

Chart1
| SPECIES | BRAZILIAN<br>POPULAR<br>NAME | REASON FOR<br>HOSPITALIZATION | CITY FROM | RT-qPCR<br>RV | RT-PCR<br>RV | PBV |
| --- | --- | --- | --- | --- | --- | --- |
| *Turdus rufiventris* | Sabiá-laranjeira | Diarrhea | Castanhal | 39.1 | G1 | Neg |
| *Turdus sp.* | Sabiá | Diarrhea | Paragominas | 34.8 | G3 | Neg |
| *Serinus canaries* | Canário-do-reino | Pododermatitis, tachypnea and diarrhea | Castanhal | 38.2 | G3 | Neg |
| *Sporophila angolensis* | Curió | Diarrhea | Paragominas | 38.5 | G6 | Neg |
| *Amazona amazonica* | Papagaio-do-mangue/curica | Lameness, limb atrophy and diarrhea | Inhangapi | Neg | G3 | Neg |
| *Heterospizias meridionalis* | Gavião-caboclo | Humeral fracture and diarrhea | Castanhal | 38.5 | T2/G3 | Neg |
| *Amazona amazonica* | Papagaio-do-mangue/curica | Respiratory, cardiovascular and ocular systems injury | Santa Izabel | 33.0 | RVA/<br>RVD/<br>RVF | Neg |
| *Ramphastus sp* | Tucano | Diarrhea | Castanhal | 38.0 | G3 | G1 |
| *Nymphicus hollandicus* | Calopsita | Apathy, diarrhea and depression | Capanema | 39.0 | G3 | Neg |

**Fig 1.**
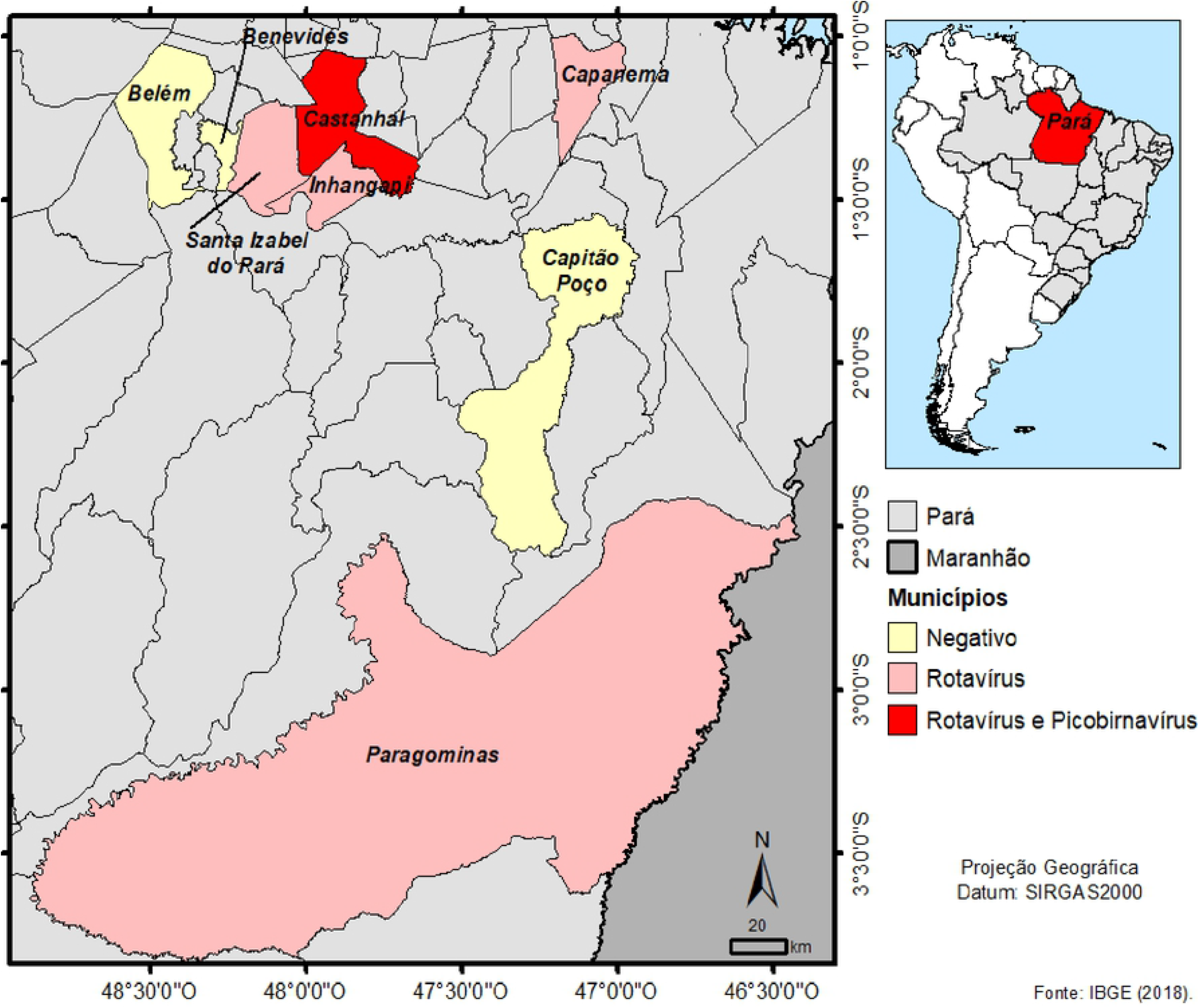
Geographic distribution of positive cases of RV and PBV in wild birds in the state of Pará.

The positive samples from RT-qPCR were amplified by RT-PCR also for the NSP3 gene and one sample (1/9) was generated with specific amplicon of 1078 bp. This sample came from an adult Savanna Hawk (*Heterospizias meridionalis*) of undetermined gender, which belonged to the T2 genotype of RVA, revealing 100% nucleotide homology with a human strain detected in Belgium in 2009 (JF460831), as shown in **Fig 2**.

**Fig 2.**
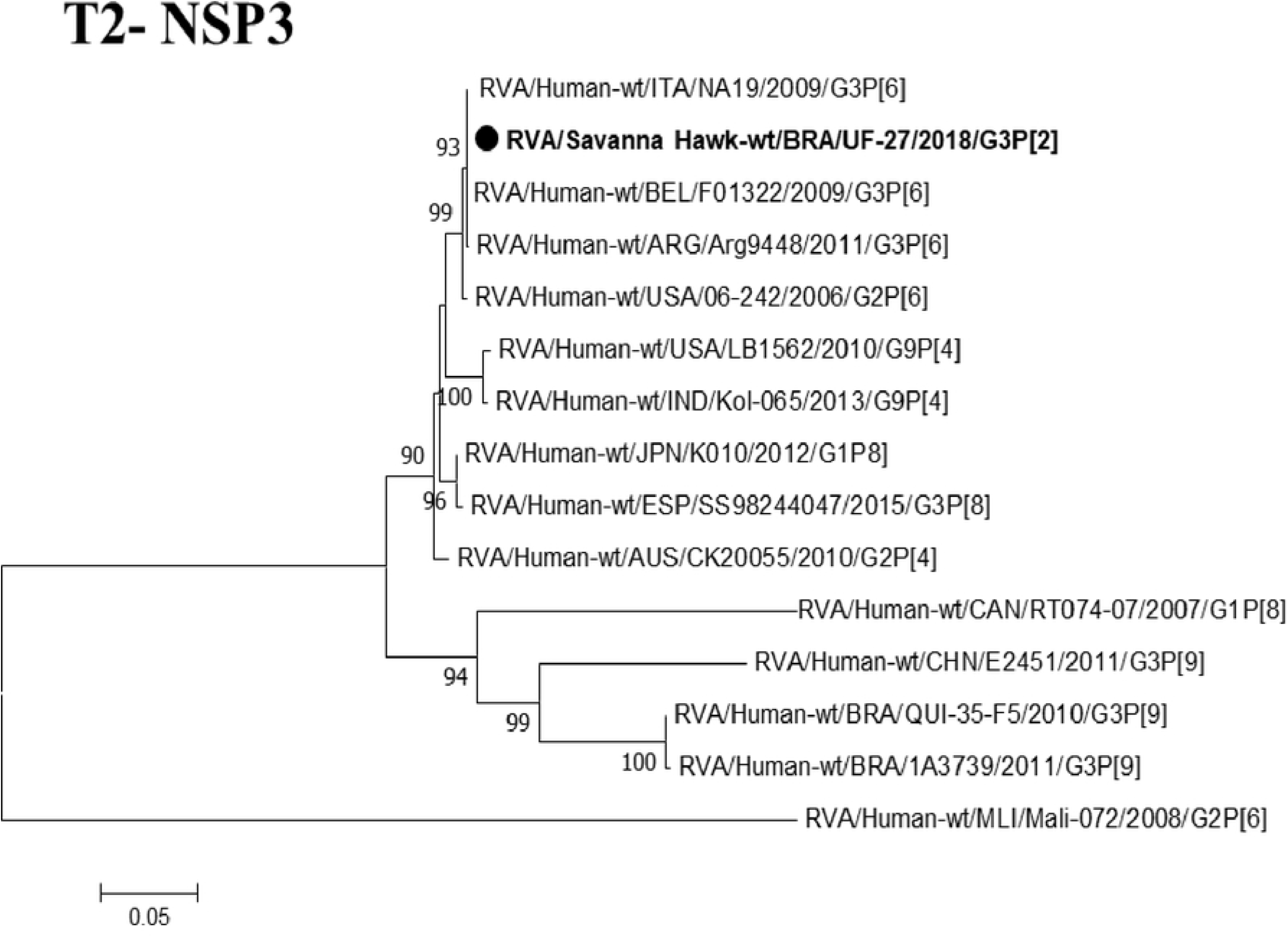
Phylogenetic tree based on the alignment of RVA NSP3 gene sequences. **Legend:** The symbol • represents the sequences of this study, and the black color depicts the birds. The numbers next to the nodes indicate bootstrap values > 70.

For the VP7 gene, it was possible to obtain the sequence of 88.9% (8/9) of the samples, generating a 193 bp amplicon (**Fig 3**). Among the eight positive samples for RVA, one diarrheic sample from Rufous-bellied Thrush (*Turdus rufiventris*) collected in Castanhal reported G1 genotype with similarity of 98% to the Russian strain (KT000090) collected in 2010. In addition, one diarrheic sample from Chestnut-bellied Seed-Finch (*Sporophila angolensis*) from Paragominas showed the human G6 genotype with similarity of 95% to an African human strain (LC026103) collected in Ghana in 2012.

**Fig 3.**
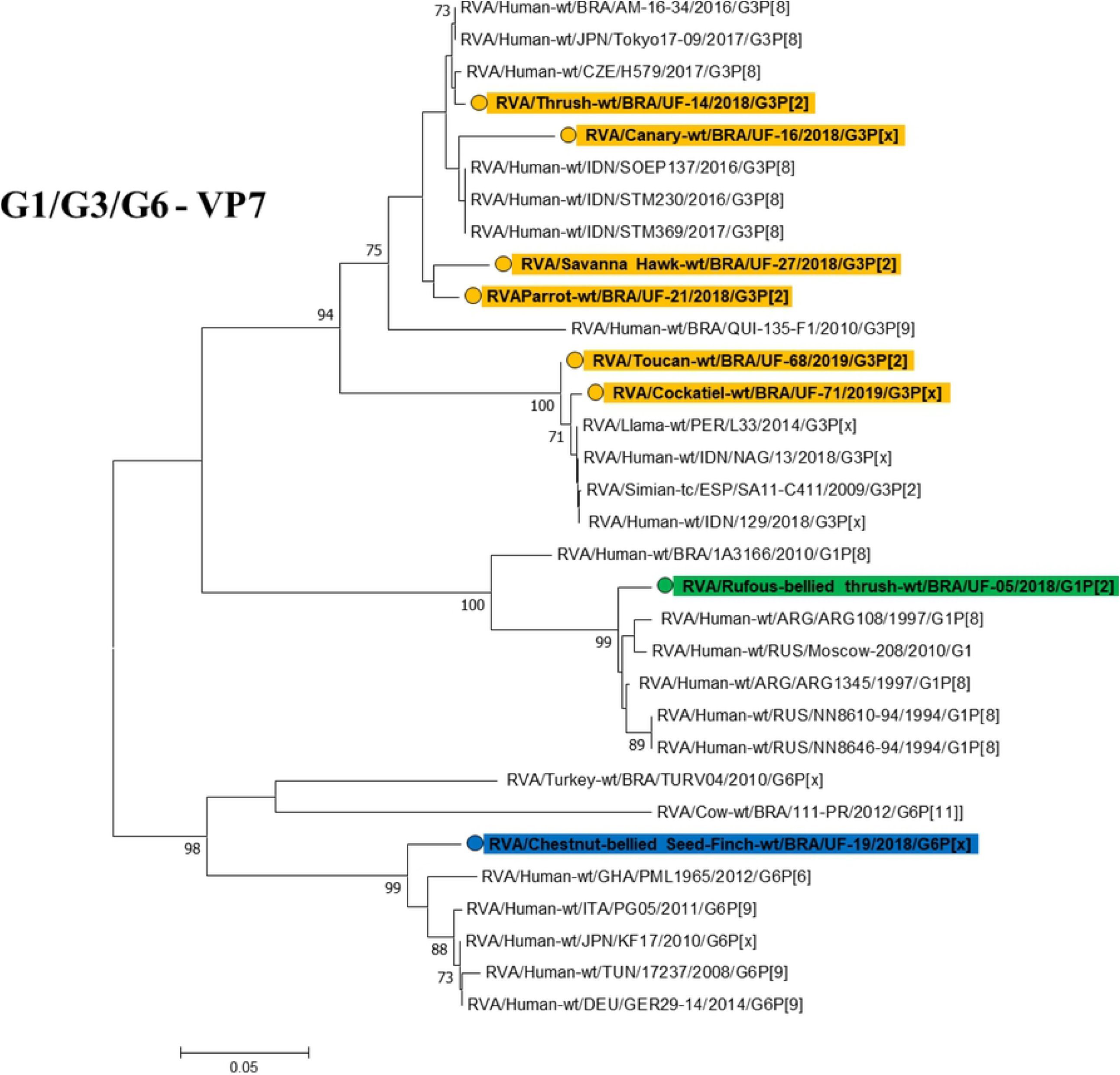
Phylogenetic tree based on RVA VP7 gene sequence alignment. **Legend:** The symbol ◯ represents the RVA VP7 gene sequences of this study, and the orange symbol • depicts the G3 genotype, the green symbol • the G1 genotype and the blue symbol • the G6 genotype. The numbers next to the nodes indicate bootstrap values > 70.

Six samples indicated the equine-like G3 genotype, with two samples belonging to an Orange-winged Parrot (*Amazona amazonica*) and Savanna Hawk (*Heterospizias meridionalis*) collected in 2018 and grouped in the same clade. They showed nucleotide similarity of 99% and 98% with human strains previously detected in the Amazon Region in 2016 (KX469402) and Japan in 2017 (LC477351, LC477356), respectively. The canary sample (*Serinus canaria*), also collected in 2018, was grouped in a different clade, reporting a similarity of 96% with human strains (LC434540) circulating in Indonesia in 2017. The canary and the Savanna Hawk were obtained from Castanhal and the Orange-winged Parrot from Inhangapi. All birds showed clinical signs of diarrhea.

The diarrhea sample from a thrush (*Turdus sp*.) came from Paragominas and formed a clade separately from the other samples of the G3 genotype, demonstrating a 99% nucleotide association with a circulating human strain in Czech Republic (MK690517) in 2017. The other two diarrheic samples belong to a toucan (*Ramphastus sp*.) and a cockatiel (*Nymphicus hollandicus*) and were obtained from Castanhal and Capanema, respectively. These specimens were grouped in the same clade with nucleotide homology of 99% for cockatiel and 100% for toucan when compared to the strains reported in llamas (KY972039), collected in Peru in 2014.

Regarding the VP4 gene, it was possible to obtain the sequence of 55.5% (5/9) of the samples, producing an amplicon with 214 bp. These specimens came from one thrush (*Turdus sp*.), one toucan (*Rhamphastus sp*.), one Orange-winged Parrot (*Amazona amazonica*), one Savanna Hawk (*Heterospizias meridionalis*) and one Rufous-bellied Thrush (*Turdus rufiventris*), which presented the P[2] genotype with 99% similarity to the human strains MH769713 and MH769713 detected in India in 2018, as shown in **Fig 4**.

**Fig 4.**
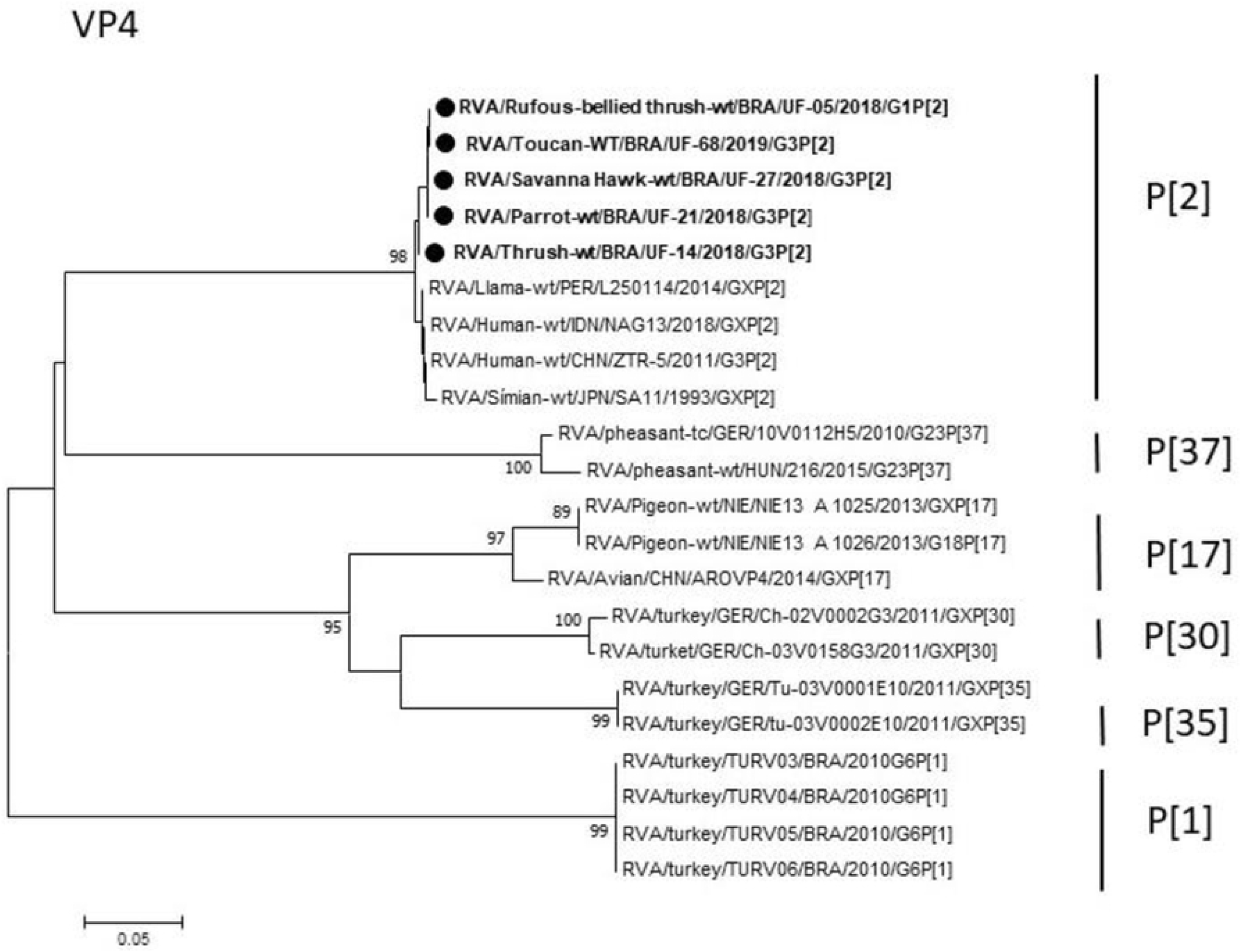
Phylogenetic tree based on the sequence alignment of the avian RVA VP4 gene. **Legend:** The symbol • represents the sequences of this study, and the black color depicts the birds. The numbers next to the nodes indicate bootstrap values > 70.

In regard to avian rotaviruses (RVA, RVD, RVF and RVG), one sample was amplified concomitantly for RVA, RVD and RVF (1/23) and produced amplicons of 642 bp, 742 bp and 846 bp for the NSP4 genes (**Fig 5**) and VP6 (**Fig 6**), respectively. The sample was obtained from an Orange-winged Parrot (*Amazona amazonica*) from Santa Izabel, which showed no clinical signs of diarrhea. The specimen reported 100% similarity with the reference strains for RVA (JN374839), RVD (KJ101587) and RVF (KP824808) circulating in the Amazon Region. No positivity was recorded for RVG.

**Fig 5.**
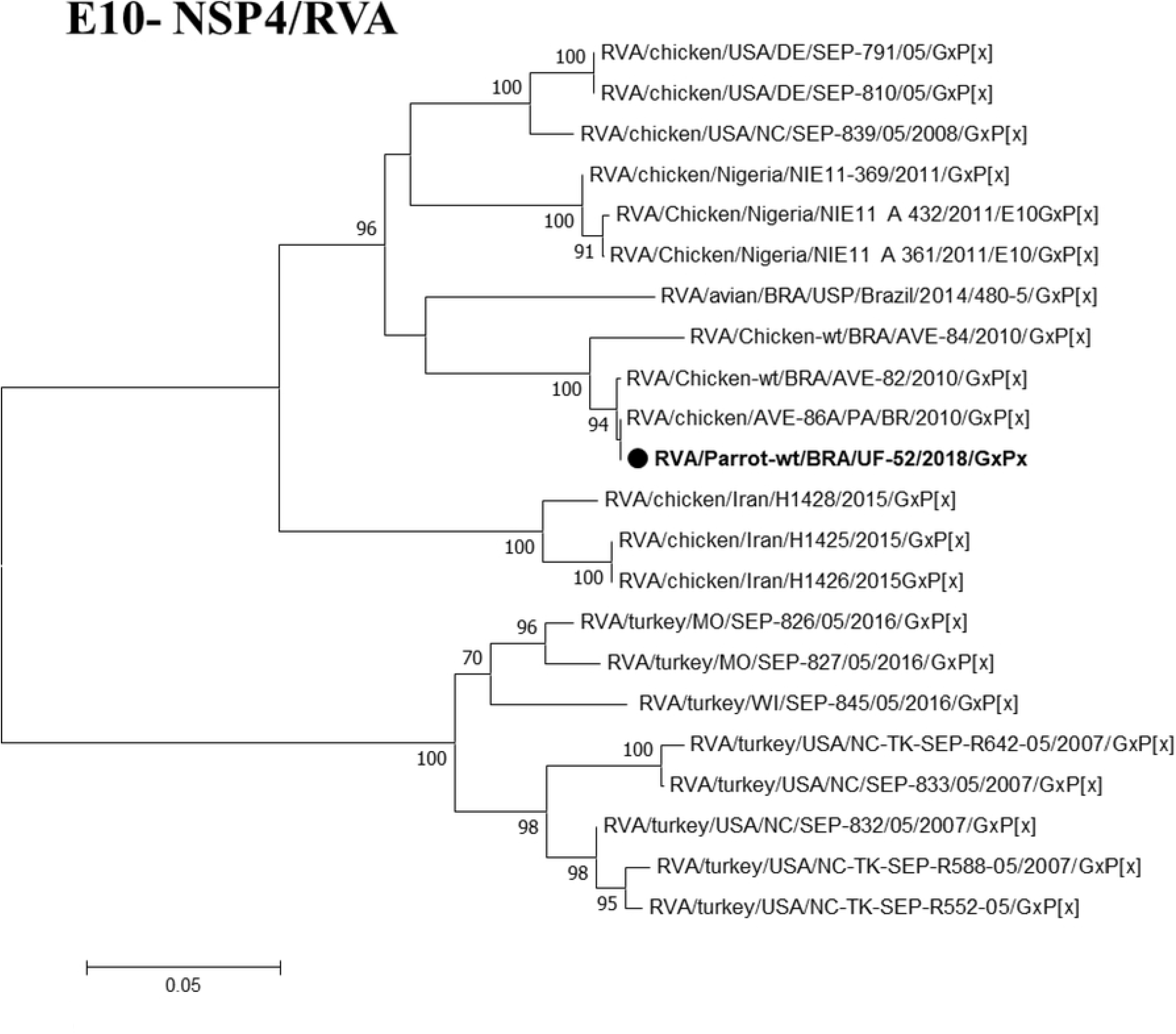
Phylogenetic tree based on the alignment of avian RVA NSP4 gene sequences. **Legend:** The symbol • represents the sequences of this study, and the black color depicts the birds. The numbers next to the nodes indicate bootstrap values > 70.

**Fig 6.**
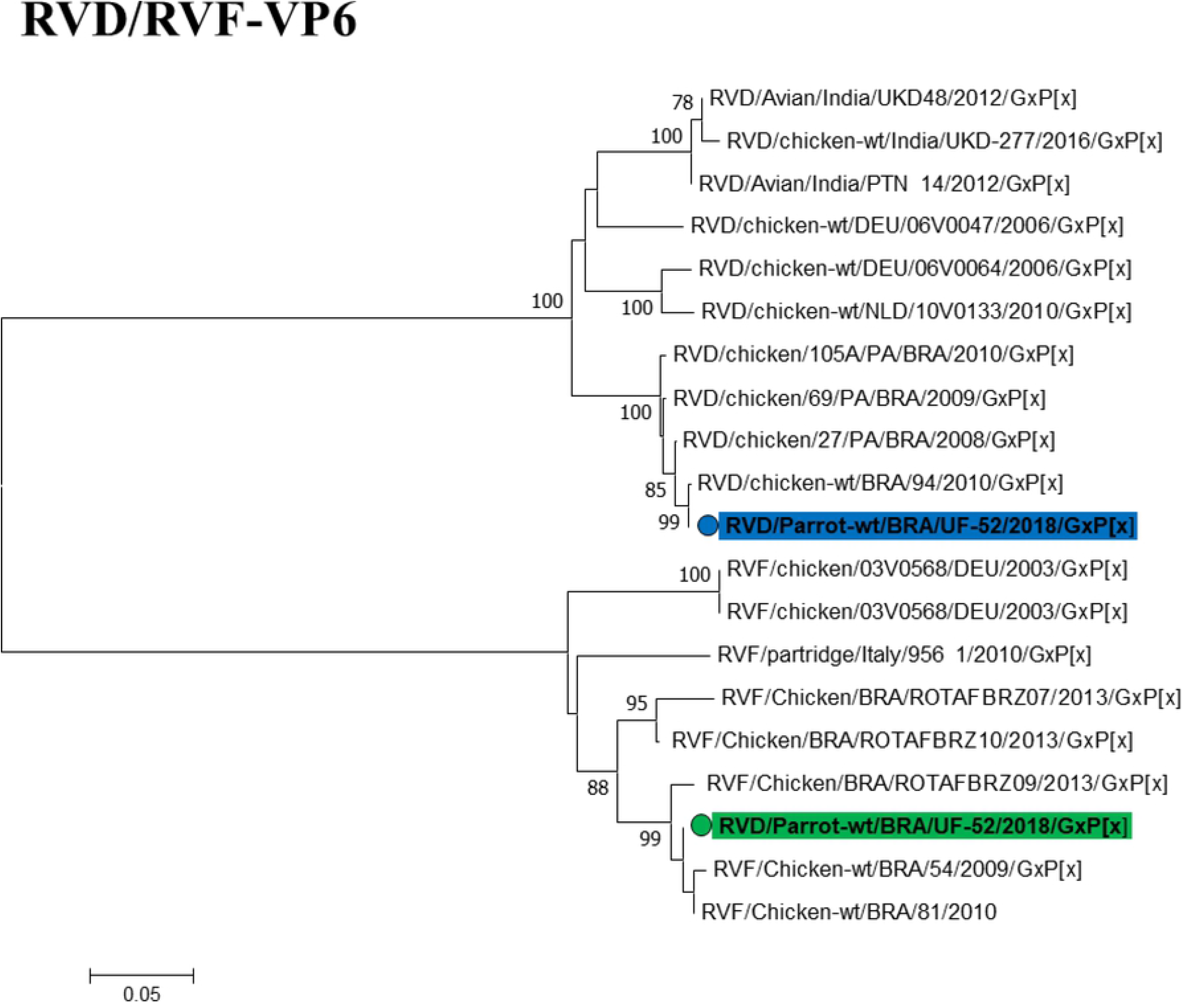
Phylogenetic tree based on the sequence alignment of the VP6 gene of RVD and avian RVF. **Legend:** The symbol ○ represents the VP6 gene sequences of rotavírus in this study, and the blue symbol • depicts the RVD, and the green symbol • the RVF. The numbers next to the nodes indicate bootstrap values > 70.

Considering PBV, among the 23 samples tested, one amplified for the RdRp gene (1/23) yielded an amplicon of 580 bp. This specimen came from a toucan (*Ramphastus sp*) from Castanhal with sign of diarrhea. The sample was grouped in Genogroup I of PBV, showing 97% nucleotide similarity with the reference strain MG846412, which was reported in Rio Grande do Sul in 2015, and 98% with KC865821 circulating in the Amazon Region in 2010, as shown in **Fig 7**.

**Fig 7.**
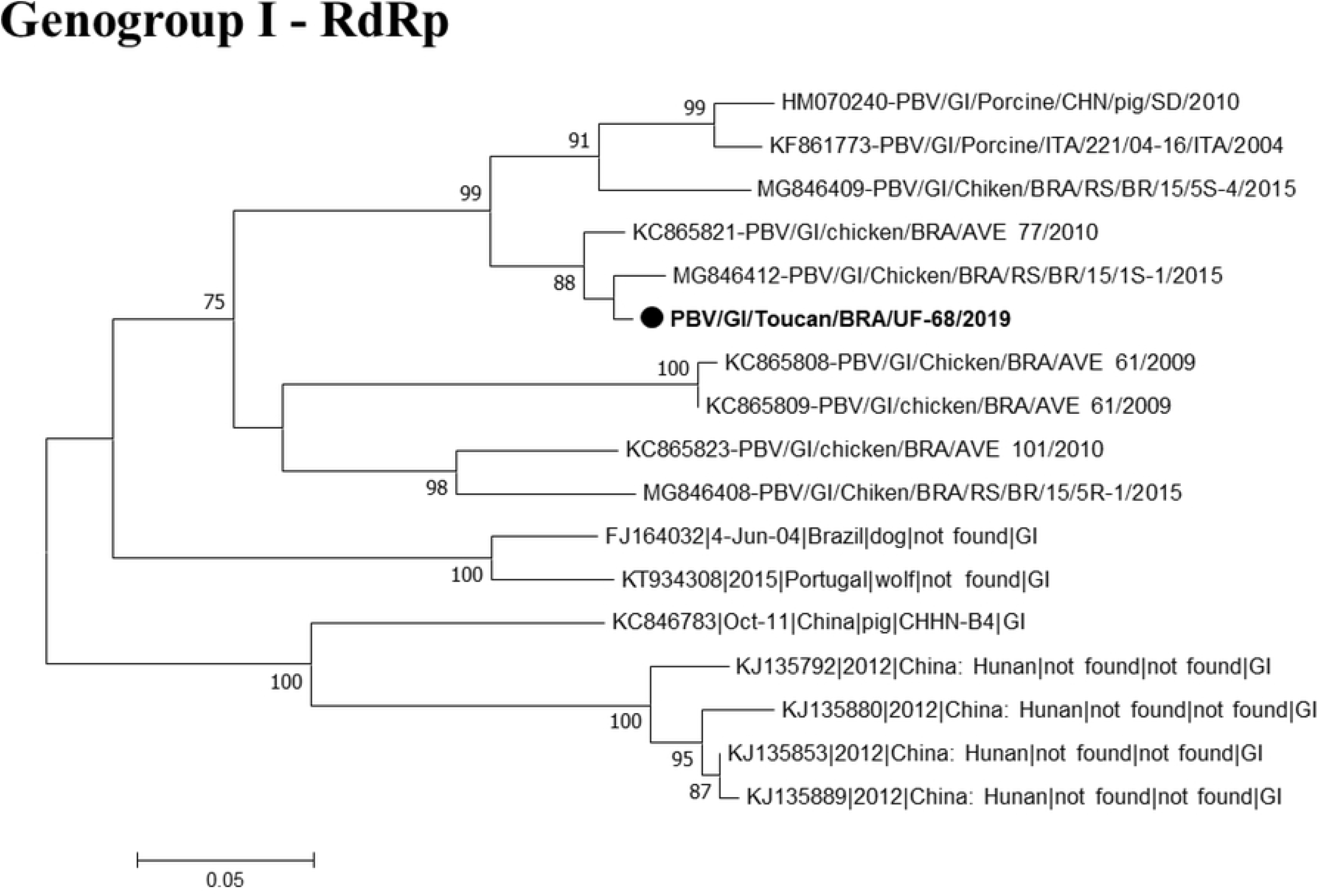
Phylogenetic tree based on sequence alignment of the PBV RdRp gene. **Legend:** The symbol • represents the sequences of this study, and the black color depicts the birds. The numbers next to the nodes indicate bootstrap values > 70.

## DISCUSSION

In the present study, RVA was detected circulating in 39.1% of the samples (9/23), while RVD, RVF and PBV were detected in 4.5% of wild birds in the Amazon avifauna. Notably, the results are contrasting with the findings of Guerreiro et al. [25] that reported negative results for the 23 fecal samples of migratory birds for rotavirus and PBV using the same primers of this study. Besides that, the low sensitivity of PAGE was recorded and may be justified due to the low viral load excreted from birds, corroborating the data from Masachessi et al. [11], Guerreiro et al. [25], Fregolente et al. [26] and Barros et al. [27], which displayed no electrophoretic profile in the positive samples.

The higher CT-values in RT-qPCR oscillated from 33 to 39.1 (mean = 37.38), hence indicating the presence of low viral load of RVA in wild specimens. Remarkably, Barros et al. [27] demonstrated that RT-qPCR assay is an efficient tool to detect RVA in specimens with low viral load, and in this study, it was further possible to perform molecular detection and characterization for NSP3, VP7 and VP4 genes of RVA, with greater recovery of RVA sequences. This occurred probably due to the use of the method for nucleic acid extraction described by Boom et al. [12], in contrast of Barros et al. [27], who used TRIzol and characterized 1.25% (8/648) of the samples only for the VP4 gene. In fact, TRIzol has the capacity to extract RNA with high integrity. However, chloroform residues can consume the RT-qPCR reagents and may cause amplification failure, hence justifying the low rate of molecular characterization reported by Barros et al. [27]. The nucleotide sequences showed more similarity to the human strains owing to the scarcity of wild birds origin nucleotide sequences [28], whereas the genotypes circulating in birds are T4 and T8 for NSP3 [17, 29-32], G5 [32], G6 [33], G7 [34], G8 [35], G10 [33], G11 [32], G17 [32], G18 [35], G19 [35], G22 [36], G23 [29] for VP7 gene [30, 33-37], and P[1] [37], P[17] [35], P[30] [17], P[31] [35], P[35] [28] and P[37] [29], P[38] [28] for VP4 gene [17, 29, 40, 36, 38].

The T2 genotype of the NSP3 gene was identified in a Savanna Hawk (*Heterospizias meridionalis*) and possessed similarity with T2 of human origin strains. Importantly, this is the first report involving the circulation of T2 genotype in wild birds. Once this sample belongs to a rescue bird and shows similarity of 100% with a Belgian human strain detected in 2009, we suggest interspecies transmission between humans and Savanna Hawk and the emergence of the Genotype T2 in the Amazon region, evidencing a possible circulation in wild birdlife.

Regarding the VP7 gene, the same hawk sample grouped in the unusual G3 genotype demonstrated human origin with phylogenetic proximity to a circulating strain in Japan (LC477351, LC477356), an Orange-winged Parrot (*Amazona amazonica*) sample grouped with samples that circulate in Brazil. These findings emphasize the importance of continuous monitoring of RVA in virtue of the previous report of the equine-like G3 genotype in the Amazon Region [39], originated from the selective pressure of zoonotic strains and due to the possible presence in different ecological niches.

The canary sample (*Serinus canaria*) grouped with the equine-like human sample reported in Indonesia in 2017 [40] and a sample from thrush (*Turdus sp*.) also formed a clade similar to human samples, but described in Czech Republic in 2017. The samples of toucan (*Ramphastus sp*.) and cockatiel (*Nymphicus hollandicus*) were grouped in a different clade, but showed high nucleotide homology (99% and 100%, respectively) to strains circulating in llamas [41]. This genotype is considered the most virulent, justifying the diarrheal condition evidenced by the animals grouped in the G3 genotype [40].

The RVA genotype G3 is considered the third most common genotype due to a larger spectrum of hosts and a greater potential for interspecies transmission in comparison to the human genotypes G1, G2 and G4. This genotype is circulating in wild animals, as well as in others, such as cattle, canines, horses, pigs, leporids, sheep, camelids, rodents, felines, simians, bats and also in humans [28]. The present study is the first to report the circulation of G3 genotype in wild birds and exotic species, and scientific evidence indicates the possible interspecies transmission or exposure of these animals to the same source of contamination.

In 2017, Bezerra et al. [42] reported the circulation of the G3 genotype in samples from quilombolas of the Amazon Region, which had similarity with G3 from animal origin (simian, bats, llama, equine and alpaca). Nevertheless, further studies on the genotypic constellation of the RV genomic segments on the specimens evaluated are needed in order to obtain data regarding their distribution and the possible natural reservoirs of the G3 genotype in wild fauna. A sample of Rufous-bellied Thrush (*Turdus rufiventris*) grouped into G1 genotype with similarity of 98% with a human strain KT000090, identified in Russia in 2010, was already reported in humans [43], ursids [44], sheep, cattle, llamas [45] and pigs [46]. The Chestnut-bellied Seed-Finch sample (*Sporophila angolensis*) grouped into genotype G6 has similarity of 95% with the human strain LC026103, found in West Africa in 2012, and has been reported in birds [38], cattle [46], sheep [47], pigs [48], horses [49], antelopes [50], leporids [51], doe [52] and human [53].

The birds of the order *Passeriformes* (seed-finch and thrush) represent the most illegally marketed group, corresponding to 90% of the avian traffic, due to their beautiful singing. In turn, the *Psittaciformes* (parrot) represent 6% due to their colored feathers, and other genera of birds correspond to 4% [54].

In this study, the human genotypes circulating in seed-finch and thrush can be justified owing to human contact arising from the illegal commercialization of these birds, characterizing the first record of genotypes G1 and G6 in wild birds.

Regarding the VP4, genotype P[2] is an uncommon one, rarely reported in the literature, which has been recently studied. Notwithstanding that, it has already been found in humans, simians [55, 56], llamas and alpacas [42]. In this study, all the samples grouped in the same clade showed distance from other samples detected worldwide and demonstrated more proximity to indigenous Brazilian human strains. Therefore, it is the first report of the circulation of genotype P[2] in Brazil. Recently, Rojas et al. [41] reported the unusual genotype P[2] circulating in llamas, alpacas and humans in Peru, thus revealing the zoonotic potential associated with the circulation of genotypes G1 and G3 in these animals. The findings suggest the close interaction of humans and wild animals that can result in the breaking of the barrier between species, resulting in the adaptation of RV to new hosts.

These data corroborate with Asano et al. [38] and Da Silva et al. [57] who reported bovine and pig genotypes in birds, emphasizing the interspecies transmission of RVA involved in the wild cycle on Amazon. Previous study of Luchs and Timenetsky [28] showed the prevalence of genotype G3 in wildlife. However, noted birds represent 80% of animal species illegally marketed in Brazil. This practice imposes a risk to biodiversity and affects the health of animals, which are often in precarious situations such as malnutrition, abrasions, immune weakness, overcrowding in cages and absence of sanitary space, and which may be exposed to different etiological agents. Because wild and exotic birds are considered potential reservoirs for zoonotic diseases, human and environmental health can also be influenced, unbalancing the biological cycle of several pathogens, justifying the presence of unusual genotypes in the specimens under study [54, 58].

In the Amazon Region, Luz et al. [59] and Guerreiro et al. [25] investigated rotavirus A and D in wild captive and migration birds, respectively. Accordingly, the researchers did not observe the circulation of RV and PBV after using the same primers of this study. Contrarily, Guerreiro et al. [25] designed a primer targeting the VP7 gene and obtained 1/23 positivity for avian RVA.

Barros et al. [27] reported the presence of RVA by RT-qPCR in 23.6% of poultry and wild birds circulating in the Amazon Brazilian biome. Bezerra et al. [18], Da Silva et al. [57] and Mascarenhas et al. [19] detected RVD, RVA, RVF and RVG in broiler chicken in the same cities explored in this study (Belém, Ananindeua, Inhangapi, Benevides, Castanhal, and Santa Izabel). Otherwise, in the current research, a sample from parrot (*Amazon amazonica*) amplified only for RVA NSP4 gene, RVD and RVF VP6 genes showed high similarity with RVA, RVD and RVF strains circulating in the Amazon. This animal came from Santa Izabel and had no clinical signs of diarrhea, but manifested deficiency of the cardiovascular, ocular and upper respiratory systems, rendering the bird susceptible to viral infections.

Regarding PBV, all samples were tested for both GG-I and GG-II, and the circulation of GG-I was identified in a toucan sample (*Ramphastus sp*) with diarrheic symptomatology, showing coinfection for RV and reporting a high nucleotide similarity of 97% with the *Chicken picobirnavirus* strain, detected by Lima et al. [32] in birds from Rio Grande do Sul, Brazil. Therefore, this is the first report of the RdRp PBV gene in wild birds in Brazil circulating in toucans from the Amazon region.

The absence of diarrheal symptoms resulting from PBV infections is reported in several hosts [10], which can be one of the factors interfering in the diagnosis, considering that the amount of viral particles excreted in the feces is not detectable by PAGE. Thus, due to the low viral load, no positive signal in PAGE was observed in this study, thereby suggesting that PBVs were not the primary agent for the manifestation of diarrheal conditions because these animals present coinfection for rotavirus.

However, it is suggested that the diarrheic animals that were not positive for PBV could be affected by other enteric agents (viral, fungal, bacterial, protozoan or helminthic) and likely have triggered diarrheal conditions due to stress caused by physiological disorders such as fractures, myiasis, tachycardia, tachypnea and other injuries. For those animals positive for RV with no signs of diarrhea, we suggested the low viral load in the samples, which may explain the absence of any manifested symptoms (observed in the CT’s obtained in the RT-qPCR).

Menes and Simonian [60] interviewed street market merchants of Bragança, Cametá, Capanema, Castanhal, Paragominas, Santarém and Tucuruí cities (state of Pará) regarding the clandestine commercialization of wild animals, and found that Castanhal exhibited the highest commercialization of these species. The data corroborate with our study considering the high prevalence of RVs and PBVs in Castanhal and also demonstrate that this illegal activity could interfere in the transmission of viral agents, including RV and PBV, between ecosystems and their genetic diversity.

Thus, Barros et al. [27], Guerreiro et al. [25], Luz et al. [59], Da Silva et al. [57], Mascarenhas et al. [19] and Bezerra et al. [18] corroborate the present study. The literature reports suggest that the Metropolitan regions of Belém and Northeast of the state of Pará concentrate the highest rates of anthropic pressure in the Legal Amazon, thus unveiling that the heterogeneity of RV and PBV in birds is co-circulating in urban, rural and wild ecosystems.

## CONCLUSION

Epidemiological data on the dynamics of enteric viruses in wildlife of the Amazon region are still scarce. This study is a pioneer in reporting the human and animal genotypes circulating in the Amazonian urban habitat. Therefore, additional evidence in wild birds and exotic species is required in order to provide a comprehensive understanding of the biological cycle of rotavirus and picobirnavirus in these animals. Further, the interactions with avian species and dispersal among human and other animals, as well as the identification of the predominant strains to determine the role of these birds in the epidemiology of the disease and to develop prophylactic measures are of utmost importance.

## ACKNOWLEDGMENTS

The authors thank the technical support of Arbovirology Molecular Biology laboratory and Rotavirus Laboratory of the Evandro Chagas Institute. We are grateful to the partnership of the Veterinarian Hospital of UFPA (HOVET-UFPA), Laboratory of Geoprocessing (LABGEO) and Dr. Yashpaul Malik from Indian Veterinary Research Institute. Thanks are also given to the National Development Agency Council for Scientific and Technological Development (CNPq), Coordination of Improvement of Higher Education Personnel (CAPES) and Amazonia Paraense Foundation for Research Support (FAPESPA) for financial support.

## Reference

1. Piacentini VDQ, Aleixo A, Agne CE, Maurício GN, Pacheco JF, Bravo GAC. Annotated checklist of the birds of Brazil by the Brazilian Ornithological Records Committee/Lista comentada das aves do Brasil pelo Comitê Brasileiro de Registros Ornitológicos. Revista Brasileira de Ornitologia, v. 23, n. 2, p. 91–298, 2015.

2. BirdLife International Country profile: Brazil [Internet]. 2019 [cited 2019 October 30] Avaible from http://www.birdlife.org/datazone/country/brazil.

3. INPE – INSTITUTO NACIONAL DE PESQUISAS ESPACIAIS [Internet]. 2018 [cited 2019 October 30]. Taxas anuais de desmatamento na Amazônia Legal Brasileira (AMZ). Avaible from: http://www.obt.inpe.br/prodes/dashboard/prodes-rates.html.

4. Martella V, Bάnyai K, Matthijnssens J, Buonavoglia C, Ciarlet M. Zoonotic aspects of rotaviruses. Vet Microbiol, v. 140, n. 3 - 4, p. 246 – 255, 2010.

5. Magle SB, Hunt VM, Vernon M, Croocks KR. Urban wildlife research: past, present, and future. Biological Conservation, v. 155, p. 23–32, 2012.

6. Estes MK, Greenberg HB. Rotaviruses In: sKNIPE DM, HOWLEY P. M. Fields Virology, 5 ed. Philadelphia: Lippincott Williams & Wilkins. p. 1347 – 1401. 2013.

7. Phan TG, Leutenegger CM, Chan R, Delwart E. Rotavirus I in feces of a cat with diarrhea. Virus genes, v. 53, n. 3, p. 487–490, 2017.

8. Delmas B, Attoui H, Ghosh S, Malik YS, Mundt E, Vakharia VN. ICTV virus taxonomy profile: Picobirnaviridae. UMBC Faculty Collection, 2018.

9. Malik, Y. S., Ghosh, S. Etymologia: Picobirnavirus. Emerging Infectious Diseases, 26(1), 89, 2019.

10. Ganesh B, Masachessi G, Mladenova Z. Animal picobirnavirus. Virus disease, v. 25, n. 2, p. 223–238, 2014.

11. Masachessi G, Martínez LC, Giordano MO, Barril PA, Isa BM, Ferreyra L, et al. Picobirnavirus (PBV) natural hosts in captivity and virus excretion pattern in infected animals. Archives of virology, v. 152, n. 5, p. 989, 2007.

12. Boom RCJA, Sol CJ, Salimans MM, Jansen CL, Wertheim-Van Dillen Pm, et al. Rapid and simple method for purification of nucleic acids. Journal of clinical microbiology, v. 28, n. 3, p. 495–503, 1990.

13. Pereira HG, Azeredto RS, Sutmoller F, Leite JP, Farias VD, Barth OM, et al. Comparasion of polyacrylamide gel electrophoresis (PAGE), immuno-electron microscopy (IEM) and enzyme immunoassay (EIA) for the rapid diagnosis of rotavirus infection in children. Memórias do Instituto Oswaldo Cruz, v. 78, n. 4, p. 483–490, 1983.

14. Zeng SQ, Halkosalo A, Salminen M, Szakal ED, Puustinen L, Vesikari T. One-step quantitative RT-PCR for the detection of rotavirus in acute gastroenteritis. Journal of virological methods, v. 153, n. 2, p. 238–240, 2008.

15. Matthijnssens J, Rahman M, Martella V, Xuelei Y, De Vos S, De Leener K, Cialert M, et al. Full genomic analysis of human rotavirus strain B4106 and lapine rotavirus strain 30/96 provides evidence for interspecies transmission. Journal of virology, v. 80, n. 8, p. 3801–3810, 2006.

16. Mijatovic-Rustempasic S, Esona MD, Williams AL, Bowen MD. Sensitive and specific nested PCR assay for detection of rotavirus A in samples with a low viral load. Journal of virological methods, 236, 41–46, 2016

17. Trojnar E, Otto P, Johne R. The first complete genome sequence of a chicken group A rotavirus indicates independent evolution of mammalian and avian strains. Virology, v. 386, n. 2, p. 325–333, 2009.

18. Bezerra DAM, Da Silva RR, Kaiano JHL, Silvestre RVD, De Souza OD, Linhares AC, et al. Detection of avian group D rotavirus using the polymerase chain reaction for the VP6 gene. Journal of virological methods, v. 185, n. 2, p. 189–192, 2012.

19. Mascarenhas J D, Bezerra DA, Silva RR, Silva MJ, Júnior ECS, Soares LS. Detection of the VP6 gene of group F and G rotaviruses in broiler chicken fecal samples from the Amazon region of Brazil. Archives of virology, v. 161, n. 8, p. 2263–2268, 2016.

20. Malik YS, Sircar S, Dhama K, Singh R, Ghosh S, Bάnyai K, et al. Molecular epidemiology and characterization of picobirnaviruses in small ruminant populations in India. Infection, Genetics and Evolution, v. 63, p. 39–42, 2018.

21. Sanger F, Nicklen S, Coulson AR. DNA sequencing with chain-terminating inhibitors. Proceedings of the national academy of sciences, v. 74, n. 12, p. 5463–5467, 1977.

22. Kearse M, Moir R, Wilson A, Stones-Havas S, Cheung M, Sturrock S, et al. Geneious Basic: an integrated and extendable desktop software platform for the organization and analysis of sequence data. Bioinformatics, v. 28, n. 12, p. 1647–1649, 2012.

23. Tamura K, Stecher G, Peterson D, Filipski A, Kumar S. MEGA6: molecular evolutionary genetics analysis version 6.0. Molecular biology and evolution, 30(12), 2725–2729, 2013.

24. Kimura, MOTOO. A simple method for estimating evolutionary rates of base substitutions through comparative studies of nucleotide sequences. Journal of molecular evolution, v. 16, n. 2, p. 111–120, 1980.2.

25. Guerreiro AN, Moraes CCG, Marinho ANR, Barros BCV, Bezerra DAM, Bandeira RR, et al. Investigation of Enteric Viruses in the Feces of Neotropical Migratory Birds Captured on the Coast of the State of Pará, Brazil. Brazilian Journal of Poultry Science, v. 20, n. 1, p. 161–168, 2018.

26. Fregolente MCD, De Castro-Dias E, Martins SS, Spilki FR, Allegretti SM, Gatti MSV. Molecular characterization of picobirnaviruses from new hosts. Virus research, v. 143, n. 1, p. 134–136, 2009.

27. Barros BDCV, Chagas EN, Bezerra LW, Ribeiro LG, Júnior JWBD, Pereira D, Junior ETP, Silva JR, Bezerra DAM, Bandeira RS, Pinheiro, HHC, Guerra SFS, Guimarães RJPS, Mascarenhas JDP. Rotavirus A in wild and domestic animals from areas with environmental degradation in the Brazilian Amazon. PloS one, v. 13, n. 12, p. e0209005, 2018.

28. Luchs A, Timenetsky MCST. Gastroenterite por rotavírus do grupo A: era pós-vacinal, genótipos e transmissão zoonótica. Einstein, v. 14, n. 2, 2016.

29. Fujii Y, Mitake H, Yamada D, Nagai M, Okadera K, Ito N, et al. Genome sequences of rotavirus A strains Ty-1 and Ty-3, isolated from turkeys in Ireland in 1979. Genome Announc., 4(1), e01565–15. 2016.

30. Gάl J, Marton S, Ihάsz K, Papp H, Jakab F, Malik YS, et al. Complete genome sequence of a genotype G23P [37] pheasant rotavirus strain identified in Hungary. Genome Announc., 4(2), e00119–16. 2016.

31. Mccowan C, Crameri S, Kocak A, Shan S, Fegan M, Forshaw D et al. A novel group A rotavirus associated with acute illness and hepatic necrosis in pigeons (Columba livia), in Australia. PloS one, 13(9), e0203853. 2018.

32. Lima DA, Cibulski SP, Finkler F, Teixeira TF, Varela APM, Cerva C, et al. Faecal virome of healthy chickens reveals a large diversity of the eukaryote viral community, including novel circular ssDNA viruses. Journal of General Virology, 98(4), 690–703. 2017.

33. Beserra LAR, Barbosa BRP, Bernardes NTCG, Brandão PE, Gregori F. Occurrence and characterization of rotavirus A in broilers, layers, and broiler breeders from Brazilian poultry farms. Avian diseases, v. 58, n. 1, p. 153–157, 2013.

34. Silva LC, Sanches AA, Gregori F, Brandão PE, Alfieri AA, Headley SA, et al. First description of group A rotavirus from fecal samples of ostriches (Struthio camelus). Research in veterinary science, v. 93, n. 2, p. 1066–1069, 2012.

35. Nishikawa K, Hoshino Y, Gorziglia M. Sequence of the VP7 gene of chicken rotavirus Ch2 strain of serotype 7 rotavirus. Virology, v. 185, n. 2, p. 853–856, 1991.

36. Pauly M, Oni OO, Sausy A, Owoade AA, Adeyefa CA, Muller CP, et al. Molecular epidemiology of Avian Rotaviruses Group A and D shed by different bird species in Nigeria. Virology journal, v. 14, n. 1, p. 111, 2017.

37. Schumann T, Hotzel H, Otto P, Johne R. Evidence of interspecies transmission and reassortment among avian group A rotaviruses. Virology, v. 386, n. 2, p. 334–343, 2009.

38. Asano KM, Gregori F, Souza SP, Rotava D, Oliveira RN, Villarreal LYB, et al. Bovine rotavirus in turkeys with enteritis. Avian diseases, 55(4), 697-700. 2011.

39. Guerra S, Soares L, Lobo P, Penha Junior E, Sousa Junior E, Bezerra D, Vaz L, Linhares A, Mascarenhas J. Detection of a novel equine-like G3 rotavirus associated with acute gastroenteritis in Brazil. Journal og General of Virology. 2016.

40. Athiyyah AF, Utsumi T, Wahyuni RM, Dinana Z, Yamani LN, Soetjipto S, et al. Molecular epidemiology and clinical features of rotavirus infection among pediatric patients in East Java, Indonesia during 2015-2018: dynamic changes in rotavirus genotypes from equine-like G3 to typical human G1/G3. Frontiers in microbiology, v. 10, p. 940, 2019.

41. Rojas M, Dias HG, Gonçalves JLS, Manchego A, Rosadio R, Pezo D, et al. Genetic diversity and zoonotic potential of rotavirus A strains in the southern Andean highlands, Peru. Transboundary and emerging diseases, 2019.

42. Bezerra DA, Guerra SF, Serra AC, Fecury PC, Bandeira RS, Penha Jr ET, Linhares AC, Soares LS, Mascarenhas JD. Analysis of a genotype G3P [9] rotavirus a strain that shows evidence of multiple reassortment events between animal and human rotaviruses. Journal of medical virology, v. 89, n. 6, p. 974–981, 2017.

43. Mascarenhas JDAP, Leite JPG, Lima JC, Heinemann MB, Oliveira DS, et al. Detection of a neonatal human rotavirus strain with VP4 and NSP4 genes of porcine origin. Journal of medical microbiology, v. 56, n. 4, p. 524–532, 2007.

44. Guo, L., Yan, Q., Yang, S., Wang, C., Chen, S., Yang, X., Hao, Z. Full genome sequence of giant panda rotavirus strain CH-1. Genome Announc., v. 1, n. 1, p. e00241–12, 2013.

45. Theuns S, Heylen E, Zeller M, Roukaerts ID, Desmarets LM, Van Ranst M, et al. Complete genome characterization of recent and ancient Belgian pig group A rotaviruses and assessment of their evolutionary relationship with human rotaviruses. Journal of virology, v. 89, n. 2, p. 1043–1057, 2015.

46. Jere KC, Mlera L, O’Neill HG, Peenze I, Van Dijk AA. Whole genome sequence analyses of three African bovine rotaviruses reveal that they emerged through multiple reassortment events between rotaviruses from different mammalian species. Veterinary microbiology, v. 159, n. 1-2, p. 245–250, 2012.

47. Bwogi J, Jere KC, Karamagi C, Byarugaba DK, Namuwulya P, Baliraine FN, et al. Whole genome analysis of selected human and animal rotaviruses identified in Uganda from 2012 to 2014 reveals complex genome reassortment events between human, bovine, caprine and porcine strains. PloS one, v. 12, n. 6, p. e0178855, 2017.

48. Silva FD, Espinoza LR, Tonietti PO, Barbosa BR, Gregori F. Whole-genomic analysis of 12 porcine group A rotaviruses isolated from symptomatic piglets in Brazil during the years of 2012–2013. Infection, Genetics and Evolution, v. 32, p. 239–254, 2015.

49. Ghosh S, Taniguchi K, Aida S, Ganesh B, Kobayashi N. Whole genomic analyses of equine group A rotaviruses from Japan: evidence for bovine-to-equine interspecies transmission and reassortment events. Veterinary microbiology, v. 166, n. 3-4, p. 474-485, 2013.

50. Matthijnssens J, Potgieter CA, Ciarlet M, Parreño V, Martella V, Bάnyai K, et al. Are human P [14] rotavirus strains the result of interspecies transmissions from sheep or other ungulates that belong to the mammalian order Artiodactyla?. Journal of virology, v. 83, n. 7, p. 2917–2929, 2009.

51. Schoondermark-Van De Ven E, Van Ranst M, De Bruin W, Van Den Hurk P, Zeller M, Matthijnssens J, et al. Rabbit colony infected with a bovine-like G6P [11] rotavirus strain. Veterinary microbiology, v. 166, n. 1-2, p. 154–164, 2013.

52. Jamnikar-Ciglenecki U, Kuhar U, Sturm S, Kirbis A, Racki N, Steyer A. The first detection and whole genome characterization of the G6P [15] group A rotavirus strain from roe deer. Veterinary microbiology, v. 191, p. 52–59, 2016.

53. Matthijnssens J, Ciarlet M, Heiman E, Arijs I, Delbeke T, Mcdonald SM, et al. Full genome-based classification of rotaviruses reveals a common origin between human Wa-Like and porcine rotavirus strains and human DS-1-like and bovine rotavirus strains. Journal of virology, v. 82, n. 7, p. 3204–3219, 2008.

54. Cavalcanti CDAT, Nunes VDS. O TRÁFICO DA AVIFAUNA NO NORDESTE BRASILEIRO E SUAS CONSEQUÊNCIAS SOCIOAMBIENTAIS. Revista De Ciência Veterinária E Saúde Pública, v. 6, n. 2, p. 405–415, 2019.

55. Mitchell DB, BOTH. Complete nucleotide sequence of the simian rotavirus SA11 VP4 gene. Nucleic acids research, v. 17, n. 5, p. 2122, 1989.

56. Taniguchi K, Nishikawa K, Kobayashi N, Urasawa T, Wu H, Gorziglia M, et al. Differences in plaque size and VP4 sequence found in SA11 virus clones having simian authentic VP4. Virology, v. 198, n. 1, p. 325–330, 1994.

57. Da Silva RR, Bezerra DAM, Kaiano JHL, Manno MC, De Souza Oliveira D, et al. Molecular epidemiology of avian rotavirus in fecal samples of broiler chickens in Amazon Region, Brazil, from August 2008 to May 2011. Revista Pan-Amazônica de Saúde, 4(2), 55–62. 2013.

58. Barbosa AD, Martins NRS, Magalhães DF. Zoonoses e saúde pública: riscos da proximidade humana com a fauna silvestre. Cienc Vet Trop, v. 14, p. 1–9, 2011.

59. Luz MA, Bezerra DA, Da Silva RR, Guerreiro AN, Seixas LS, Gonzaga RK, et al. Pesquisa de rotavírus em aves silvestres da região amazônica mantidas em cativeiro no estado do Pará, Brasil. Revista Brasileira de Medicina Veterinária, [s.l.], v. 36, n. 2, p. 167-173, abr/jun. 2014.

60. Menes FLS, Simonian LTL. Animais silvestres comercializados ilegalmente em algumas cidades do estado do Pará The illegal commercialization of wild animals in some cities of state of Pará, Brazil. REMEA-Revista Eletrônica do Mestrado em Educação Ambiental, v. 33, n. 1, p. 4–21, 2016.

